# Trans-generational physiological condition of embryos is conditioned by maternal thermal stress in *Octopus maya*

**DOI:** 10.1101/2022.05.12.491712

**Authors:** Omar Domínguez-Castanedo, Daniela Palomino-Cruz, Maite Mascaró, Gabriela Rodríguez-Fuentes, Oscar E. Juárez, Clara E. Galindo-Sánchez, Claudia Caamal-Monsreal, Pavel Galindo Torres, Fernando Díaz, Carlos Rosas

**Affiliations:** Consejo Nacional de Ciencia y Tecnología. Programa de Posdoctorado; Instituto Tecnológico de La Paz, Secretaría de Educación Pública, La Paz, BCS, México; Unidad de Química Sisal, Facultad de Química, Universidad Nacional Autónoma de México. Puerto de Abrigo s/n Sisal, Yucatán, México; Unidad Multidisciplinaria de Docencia e Investigación, Facultad de Ciencias, Universidad Nacional Autónoma de México, Puerto de Abrigo s/n Sisal, Yucatán, México; Departamento de Biotecnología Marina, Laboratorio de Ecofisiología, Centro de Investigación Científica y de Educación Superior de Ensenada, Baja California (CICESE)

**Keywords:** Reactive oxygen species, respiratory metabolisms, antioxidant system, life history, global climate change, thermal stress

## Abstract

Current anthropogenic global warming generates profound metabolic alterations in marine ectotherm invertebrates capable of leading a wide range of these species to extinction. The most worrying and devastating consequence may be that the effect of thermal stress overpasses the individual generations. To evaluate the transgenerational effect of thermal stress on the cephalopod *Octopus maya*, this study experimentally tests morphology, respiratory metabolism, antioxidant mechanisms, and oxidative stress indicators of the embryos incubated at two temperatures (24 and 30°C) produced by females acclimated at 24 and 30°C. The results demonstrate that, regardless of their incubation temperature, embryos from females acclimated at 30°C are smaller, show more accelerated development, and higher respiratory rates than those from females acclimated at 24°C. These embryos confirmed a greater oxidative stress degree, as well as an increased amount of soluble carbonylated proteins and catalase activity as the main enzyme during the activation development stage (even the highest in the embryos incubated at 30°C). Finally, a collapse of the antioxidant defense system was observed, measured as lower both CAT activity and GSH concentrations. Additionally, soluble carbonylated proteins reduced and GST activity increased in embryos incubated at 30°C from females maintained at high temperatures in a clear deleterious and transgenerational effect of thermal stress on this octopus species.

## I. Introduction

An increment of sea water temperature from 1 to 3 °C is expected for the next three decades (Pörtner et al., 2022). Such increment can generate profound metabolic alterations in marine ectotherm invertebrates, affecting distribution and abundance of a wide number of species (Gillooly et al., 2001; Lefevre, 2016; Madeira et al., 2016). The mitochondrial respiratory chain is one of the main producers of reactive oxygen species (ROS) (e.g., hydroxyl and superoxide radicals, singlet oxygen, etc.) and reactive nitrogen species (RNS) (e.g., nitric oxide, peroxynitrite, etc.) that are produced naturally through physiological and non-physiological processes (Feidantsis et al., 2021; Fridovich, 1986). Oxidative stress produced by ROS consists of a set of deleterious cumulative biomolecular cell damage, mainly oxidation of lipids, proteins, and nucleic acids (Storey, 1996; Hulbert et al., 2007). The cell antioxidant defense mechanisms (ANTIOX) consist in groups of enzymes (*e*.*g*. catalase, glutathione peroxidase, peroxydoxin, superoxide dismutase; Di Giulio et al., 1989) complemented by some low molecular weight (LMN) non-enzymatic molecules (*e*.*g*. glutathione, vitamins A, C, and E, bilirubin, beta-carotene, uric acid, and flavonoids; Tiwari et al., 2014). These LMN non-enzymatic molecules progressively reduce ROS into safe compounds (*e*.*g*. O_2_ and H_2_O; Halliwell, 2012) or break the autocatalytic radical chain reactions (Cadenas, 1989), respectively. However, in organisms subjected to thermal stress, ROS production can surpass the ANTIOX system, provoking cell damage and at the end, reducing survival (Pörtner, 2010; Pörtner et al., 2017; Rodríguez-Fuentes et al., 2017). The most worrying consequence of global climate change and local thermal anomalies could be that the effect of thermal stress can overpass the capacity of individual generations to neutralize ROS, with profound costs to ecology of populations and communities (Parmesan, 2006). Moreover, increasing periods of thermal stress during reproductive phases result in a stronger transgenerational response (Dupont et al., 2013; Salinas & Munch, 2012).

Therefore, predicting how populations could react to changes in the environment, and taking the transgenerational thermal stress effect into account is key to understanding the full effects of elevated temperatures (Hendry et al., 2008). Evidence suggests that thermal tolerance between generations can be enhanced through thermal preconditioning of adults to trigger an epigenetic response that could increase thermal tolerance of their offspring (Klosing et al., 2019). That process has been defined as (1) cross-generation plasticity (CGP) (when the environment experienced by parents influences offspring phenotype: F0–F1); (2) multigenerational plasticity (MGP) (when the environment experienced by previous generations is evident to the F2 and beyond: F1–F2+); and (3) carry-over effects (COE) that occur within the development (e.g. embryo to larva) (Byrne et al., 2019). Even though thermal stress in parents has been suggested as an effective mechanism for coping with temperature increases, thereby, enhancing progeny performance under thermal stress through epigenetic inheritance (Eirin-Lopez and Putnam, 2019; Fellous et al., 2015; Fellous et al., 2021). Studies made on *Mytilus californicus* mussel provide evidence for negative parental effects on offspring thermal tolerances (Waite and Sorte, 2022).

*Octopus maya* (an endemic cephalopod adapted to comparatively low environmental temperatures of the Yucatan Peninsula) has been documented as a relatively sensitive tropical species to thermal stress (Ángeles-González et al., 2020; Ángeles-González et al., 2021). When temperature was evaluated (above 27ºC) in adult *O. maya* females, the number of spawned and fertilized eggs was lower than in females maintained at 24ºC (Juárez et al., 2015). Essential genes involved in fertilization and egg-laying downregulated in both optic and oviducal glands of thermally stressed *O. maya* females (Domínguez-Estrada et al., 2022; Juárez et al., 2022). These and more recent results have indicated that high temperatures affect negatively, not only the capacity of females to produce eggs but also how sperms fertilize the eggs in the oviductal gland (Juárez et al., 2022). In experiments with male octopuses, temperatures of 28ºC or higher provoked testicular damage, reducing their paternal contribution to progeny (López-Galindo et al., 2018). These results evidence that temperatures from 28°C to 30°C deeply affected the reproductive performance of this octopus species.

Transgenerational effects of thermal stress were documented previously in this octopus species (Juárez et al., 2016) where thermally stressed females produced smaller embryos, and the metabolic rates of their juveniles were twice those produced by non-stressed females. The authors suggested that such effects could have epigenetic bases (Juárez et al., 2015; 2016). In a study conducted on *O. mimus* embryos, Olivares et al. (2019) hypothesized that negative parental influence on offspring is related to ROS transferred from females to oocytes during oogenesis, causing an ulterior overload for embryos. In this regard, during spawning, female optic glands (located in the brain) focus on maintaining balance between energy and ROS production (Ventura et al., 2022), but high temperatures drastically affect this process (Domínguez et al., 2022). Moreover, at high temperatures, the optic gland in females up-regulates the expression of genes related to stress response and production of stress molecules like corticosterone (Dominguez-Estrada et al., 2022), which may alter embryonic development (Hayward and Wingfield, 2004; Vagnerová et al., 2008).

During the reproductive phase, females use their energy reserves for parental care (Lin et al., 2019; Roumbedakis et al., 2017; Ventura-López et al., 2022). Biological processes like oxidation-reduction process, respiratory electron transport chain and precursor metabolite and energy generation are conspicuous during spawning in this octopus species (Ventura-López et al., 2022; Meza-Buendia et al., 2021). However, thermally stressed females showed a lower metabolic scope than females maintained in thermally optimal conditions. Therefore, elevated temperatures could be affecting the ability of females to channel enough energy to all events involved in reproduction (Meza-Buendia et al., 2021).

According to Blount et al. (2016), parents avoid transferring oxidant damage to the offspring by upregulating their oxidative stress during the reproductive period. In a study made on *O. mimus*, Olivares et al. postulated that under thermal stress, female octopus fail to reduce oxidative damage transfer to their embryos, provoking an additional charge of ROS that affects embryonic development (Olivares et al., 2019). Additionally, *O. vulgaris* and *O. maya* embryos showed limited abilities to neutralize ROS when maternal charge is summed to ROS produced during their growth. ROS increases as the metabolic rate reaches its maximum level at temperatures beyond the thermal optima (Repolho et al., 2014; Sánchez-García et al., 2017).

Based on the above, this research study addresses four questions to know the effect of maternal thermal stress transferred to *O. maya* embryos: (*i*) Do embryos from thermally stressed females show alterations in their development (*e*.*g*. smaller amount of yolk), regardless of their incubation temperature? (*ii*) Does maternal thermal stress before spawning affect the metabolic response of embryos (*e*.*g*. faster respiratory rates)? (*iii*) Does maternal thermal stress and/or high incubation temperature increase oxidative stress in embryos at a specific developmental stage? and (*iv*) Is the antioxidant system of embryos from thermally stressed females subjected to thermal stress during their incubation able to cope with oxidative stress? Based on these questions and the fact that females under thermal stress may not be able to regulate ROS or oxidized biomolecule transfer to their embryos, as previously suggested by the studies done in octopus (Olivares et al., 2019; Meza-Buendia et al., 2021), their embryos are expected to show deleterious morphophysiological differences due to a decreased antioxidant system because of overwhelming oxidative stress. Moreover, those incubated under thermal stress, will not be able to metabolically compensate it.

To answer these questions, this research was directed to experimentally test in the cephalopod *Octopus maya*, the effect of thermal maternal stress on embryonic morphology, metabolic rate, oxidative damage, and antioxidant defense mechanisms in embryos exposed to thermal and optimal temperatures during their development.

## 2. Material and Methods

### 2.1 Animals and experimental design

Wild mature males and females were collected on the coast of Sisal, Yucatán by the local capture method called “gareteo”, which comes from adrift (“al garete” in Spanish, Secretaría de Agricultura, Ganadería, Desarrollo Rural, Pesca y Alimentación, 2014). The octopuses were then transferred to the experimental cephalopod production unit at the Teaching and Research Multidisciplinary Unity (UMDI-UNAM) at Universidad Autónoma de México located approximately from 1 to 2 km far from the sampling zone. The octopuses were conditioned for 2 weeks in 6000-L open-air tanks, allowing them to copulate. In the tanks, animals were maintained from 26 to 28°C. During that time, brooders were fed *ad libitum* twice aday with a paste made with squid (*Dosidicus gigas*), and crab (*Callinectes spp*) bound gelatin (Tercero-Iglesias et al., 2015). Then, six females were acclimated into individual dark tanks connected to seawater recirculation system with continuous seawater flow at 24ºC (n = 3) or 30 ºC (n = 3) for 30 days. Seawater in tanks was kept in a semi-closed recirculation system coupled with a rapid-rate sand filter and 36 ± 1 part per thousand (ppt) salinity, dissolved oxygen higher than 5 mg/L, pH above 8, photoperiod of 12L/12D and a light intensity of 30 Lux in the tank surface. Seawater temperature was gradually increased by 1ºC/day until the experimental temperature (30ºC) was reached and maintained with two 1 800 Watt heaters connected to automatic temperature controllers. The temperature of 24ºC was controlled with a titanium chiller and room air conditioner. Octopus females were maintained until spawning, with fiberglass boxes that served as a nest. The animals were fed twice a day with the previously described paste (Tercero-Iglesias et al., 2015) during the conditioning period and maintained in their experimental temperatures for 30 days. Embryos of each female were obtained after eight days, just when spawning ends. After, 80 eggs per spawn (240 embryos per experimental temperature) were randomly sampled and artificially incubated at 24ºC or 30°C (Fig. 1).

**Fig. 1.**
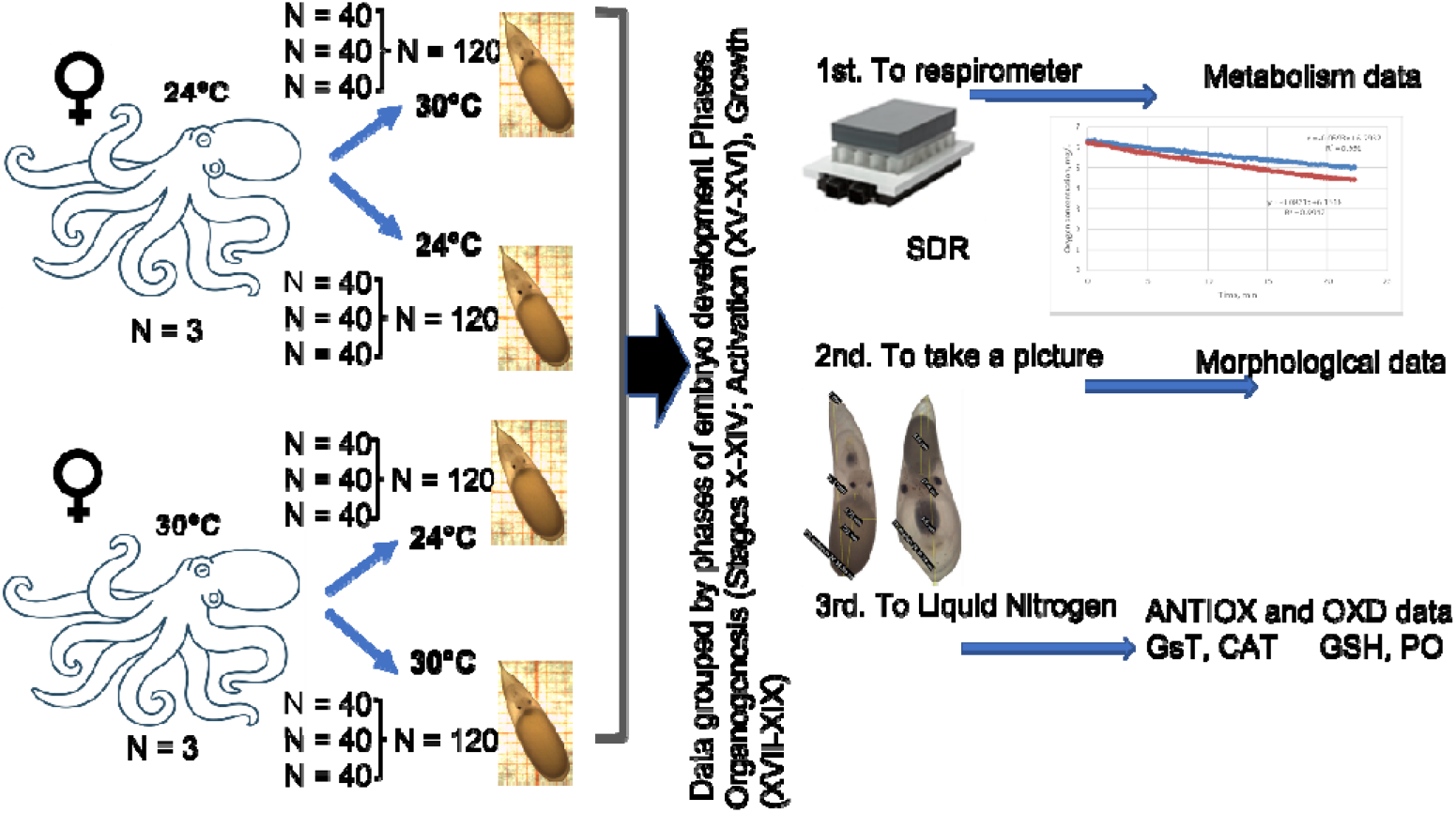
Experimental design and sample procedures to evaluate the effect of transgenerational thermal stress on the physiological condition of *Octupus maya* embryos. Please note that females were exposed for 30 days at each experimental temperature and that the embryos sampled every 5 days were used to evaluate oxygen consumption, morphological changes during development, antioxidant defense mechanisms (ANTIOX) and oxidant damage (OXD).

### 2.2 Routine oxygen consumption *in vivo*

Every five days, respiratory metabolism was measured individually: embryos were placed in micro-plate clear glass vials with integrated sensor spots (750 μL volume, Loligo Systems, Copenhagen, DK) and maintained in seawater in experimental incubation temperature (Fig. 1). Simultaneously, oxygen consumption of five control chambers (vials without an embryo) was also measured. Vials were submerged in a transparent glass container with temperature-controlled seawater maintained at 24 or 30°C. The container was placed on a Sensor Dish Reader (Presens, Regendburg, DE) that recorded oxygen concentration measurements every 15 s. Measurement time was adjusted according to oxygen concentration in the vial, which was never lower than 80% of saturation level. All measurements were graphed according to time and embryo oxygen consumption (MO_2_) calculated as:

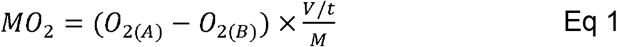

where *MO*_2_ is respiration rate (mg O_2_ h^−1^ mg WW^−1^); *O*_2(*A*)_ is initial oxygen concentration in the chamber (mg O_2_ L^−1^); *O*_2(*B*)_ is final oxygen concentration in the chamber (mg O_2_ L^−1^); *V* is water volume in the chamber minus water volume displaced by the embryo (expressed as L); *t* is time elapsed during measurement (h); and *M* is egg body mass containing the embryo (mg WW).

### 2.3 Embryonic development

Immediately after respiration rate measurements, eggs and their corresponding embryos were photographed, (Fig.1) maintaining eggs in the same experimental temperature. Morphological parameters of *O. maya* eggs (eye diameter, mm; arm length, mm; yolk length, mm; egg length, mm; egg wide, mm; and wet weight, g) were measured. Each egg was observed and photographed using Leica EZ4 HD (Wetzlar, DE) stereoscopic microscope whose software (Leica LAS EZ, Wetzlar, DE) allows identification of embryonic stage and size. The developmental stages were identified (Caamal-Monsreal et al., 2016; Naef, 1928) and grouped as organogenesis (before heart activity started; stages X to XIV), activation (when the heart starts its activity; stages XV-XVI) and growth (stages XVII-XIX). Afterwards, weight (g) of each embryo was obtained (± = 0.001g), and immediately stored in liquid nitrogen on Eppendorf tubes, and stored at −80ºC until analysis.

### 2.4 Antioxidant defense mechanisms and oxidative damage

The frozen embryos, at activation and growth phases, were individually homogenized in cold buffer 0.05M Tris pH 7.4 at 100 mg tissue/mL using a Potter-Elvehjem homogenizer. Homogenate samples used for CAT, GST and esterases were centrifuged at 10,000g for 5 min at 4°C, and the supernatant was separated for analysis. All samples were stored at −80°C until analysis; all assays were done in duplicates. Catalase (CAT) activity was measured according to Góth (1991) modified by Hadwan and Abed (2016). In this method, undecomposed H_2_O_2_ is measured with ammonium molybdate after three minutes to produce a yellowish color with maximum absorbance at 374 nm. Total glutathione (GSH) was measured with Sigma-Aldrich Glutathione Assay Kit (CS0260) (St. Louis MO, US). This kit utilizes an enzymatic recycling method with glutathione reductase (Baker et al., 1990). The GHS sulfhydryl group reacts with Ellman’
ss reagent and produces a yellow-colored compound read at 405 nm. The GST activity was determined from the reaction between reduced glutathione and 1-chloro-2.4-dinitrobenzene at 340 nm (Habig and Jakoby 1981). Proteins were analyzed in supernatant according to Bradford (1976) and used to normalize enzyme activities. To evaluate oxidative damage (OXD) caused by ROS, carbonyl groups in oxidized proteins (PO) were measured in the sampled embryos, estimating PO by using the 2,4-dinitrophenylhydrazine alkaline protocols developed by Mesquita et al. (2014) and reported in nmol/mg wet weight. For this assay, 200 µl of 2,4 dinitrophenylhydrazine (10mM in 0.5 M HCL) were incubated with 200 µl of the sample homogenate and 100 µl of NaOH (6M). Absorbance was read at 450 nm after 10 min of incubation at room temperature against a blank where an equal volume of homogenization buffer substitutes the protein solution.

For this study, two esterases – acetylcholinesterase (AChE) and carboxylesterase (CbE) – were measured to evaluate the physiological condition of embryos. AChE activity was measured using a modification of the method of Ellman et al. (1961) adapted to a microplate reader (Rodríguez-Fuentes et al., 2008). Each well contained 10 µL of the enzyme supernatant and 180 µL of 5, 5-dithiobis (2 nitrobenzoic acids) (DTNB) 0.5 mM in 0.05 M Tris buffer pH 7.4. The reaction started by adding 10 µL of acetylthiocholine iodide (final concentration 1 mM). Absorbance rate of change at 405 nm was measured for 120 s; CbE activity was measured using ρ-nitrophenyl-α-arabinofuranoside (ρNPA) substrate, as indicated by Hosokawa and Satoh. (2002) with some modifications. Each assay included 25 μL of the supernatant and 200 μL of ρNPA. The reaction was recorded for 5 min at 405 nm. SOD, AChE and CbE activities were reported per mg protein in the sample (Bradford, 1976).

### 2.5 Statistical analyses

The relationship between oxygen consumption and egg wet weight was linearized using a semi-log transformation (Ln wet weight). After that, an analysis of covariance (ANCOVA) was performed to identify statistical differences between the linear models obtained. To evaluate how the maternal thermal stress affected embryo performance (morphological changes, ANTIOX and OXD) of the embryos exposed at different temperatures, a multivariate analysis, permutational MANOVA (PERMANOVA) and Principal Coordinate Analysis (PCO) were performed using Primer v 7.0 + PERMANOVA add on. Raw data of this study were stored in ZENODO repository (Domínguez-Castanedo et al., 2022).

## 3. Results

### 3.1 Routine oxygen consumption *in vivo*

When embryo routine metabolism was related to the embryo’s wet weight, the respiratory metabolism was affected by the maternal thermal condition but not by the embryo’s thermal condition (Fig. 2). For that reason, MO2 data of embryos from each thermal female treatment (24 or 30°C) were grouped. The results obtained from the ANCOVA showed that no statistical differences were recorded between the slopes obtained from the Ln MO_2_ – Ln embryo wet weight relationship of embryos from females conditioned at 24 or 30°C (F = 2.9; DFn = 1; DFd = 242; *p* = 0.09). However, statistical differences were observed between intercepts (F = 28; DFn = 1, DFd = 243; *p* = 0.001) with lower values in embryos from females maintained at 24°C (Ln MO_2 24°C_ = Ln 1.3x - Ln 0.98; *p* □ 0.0001; F = 491; DFd = 1.16) than those obtained of embryos from females maintained at 30°C (Ln MO_2 30°C_ = Ln 1.1x - ln 1.6; *p* □ 0.0001; F = 75; DFd 1.81) (Fig. 2).

**Fig. 2.**
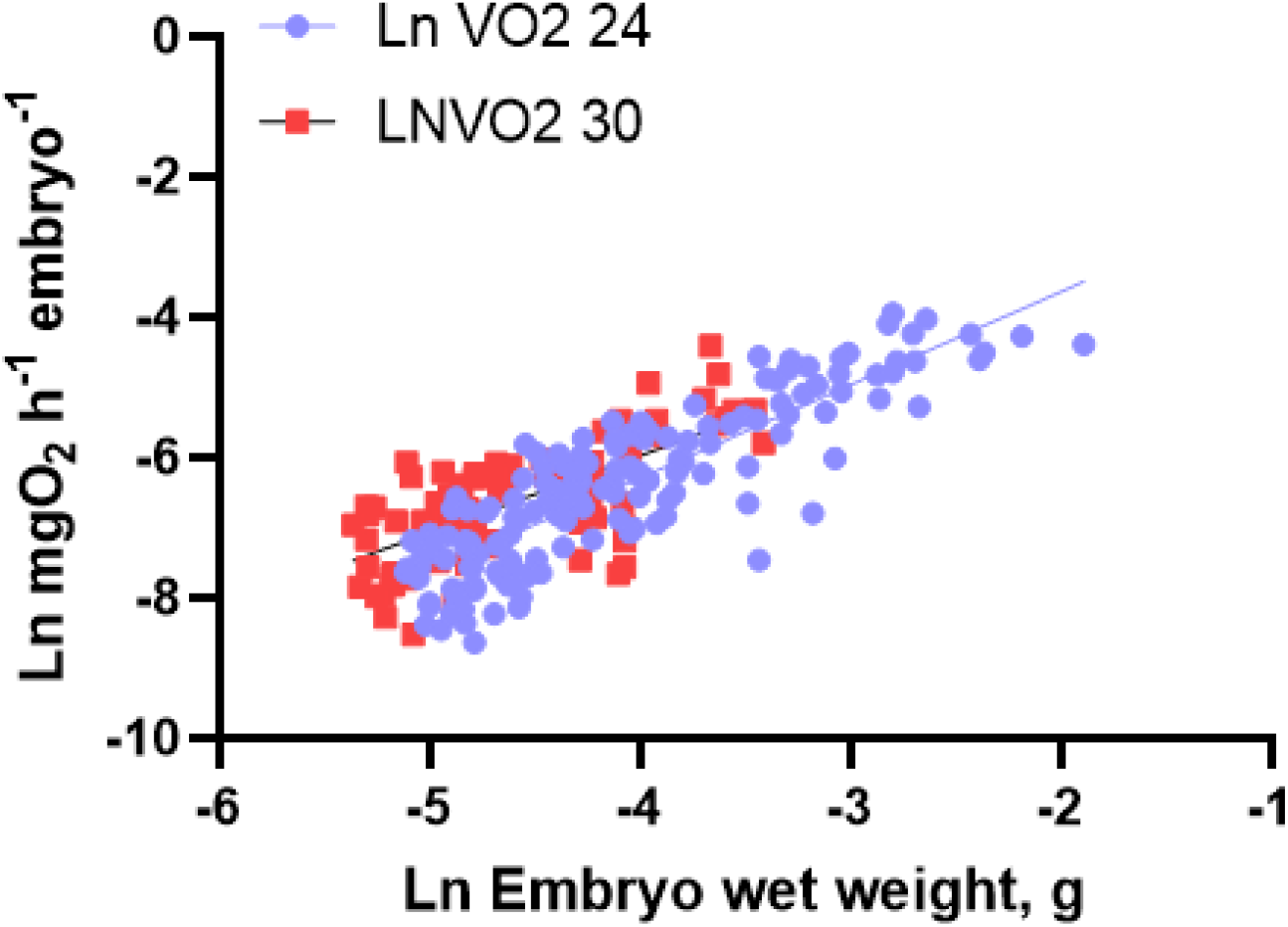
Transgenerational effect of *Octopus maya* female thermal condition (24 and 30°C) on the respiratory metabolism of embryos exposed at 24°C and 30°C. Blue dots indicate the metabolism of embryos from females conditioned at 24°C and incubated at 24 and 30°C. Red squares indicate the metabolism of embryos from females conditioned at 30°C and incubated at 24 and 30°C

### 3.2 Embryonic development

The PCO analysis indicated that the transgenerational effect of the maternal thermal condition affected the morphometric characteristics of the embryos of the next generation. Embryos from females maintained at 24°C showed larger yolks than those of embryos from females maintained at 30°C (Fig. 3).

**Fig. 3.**
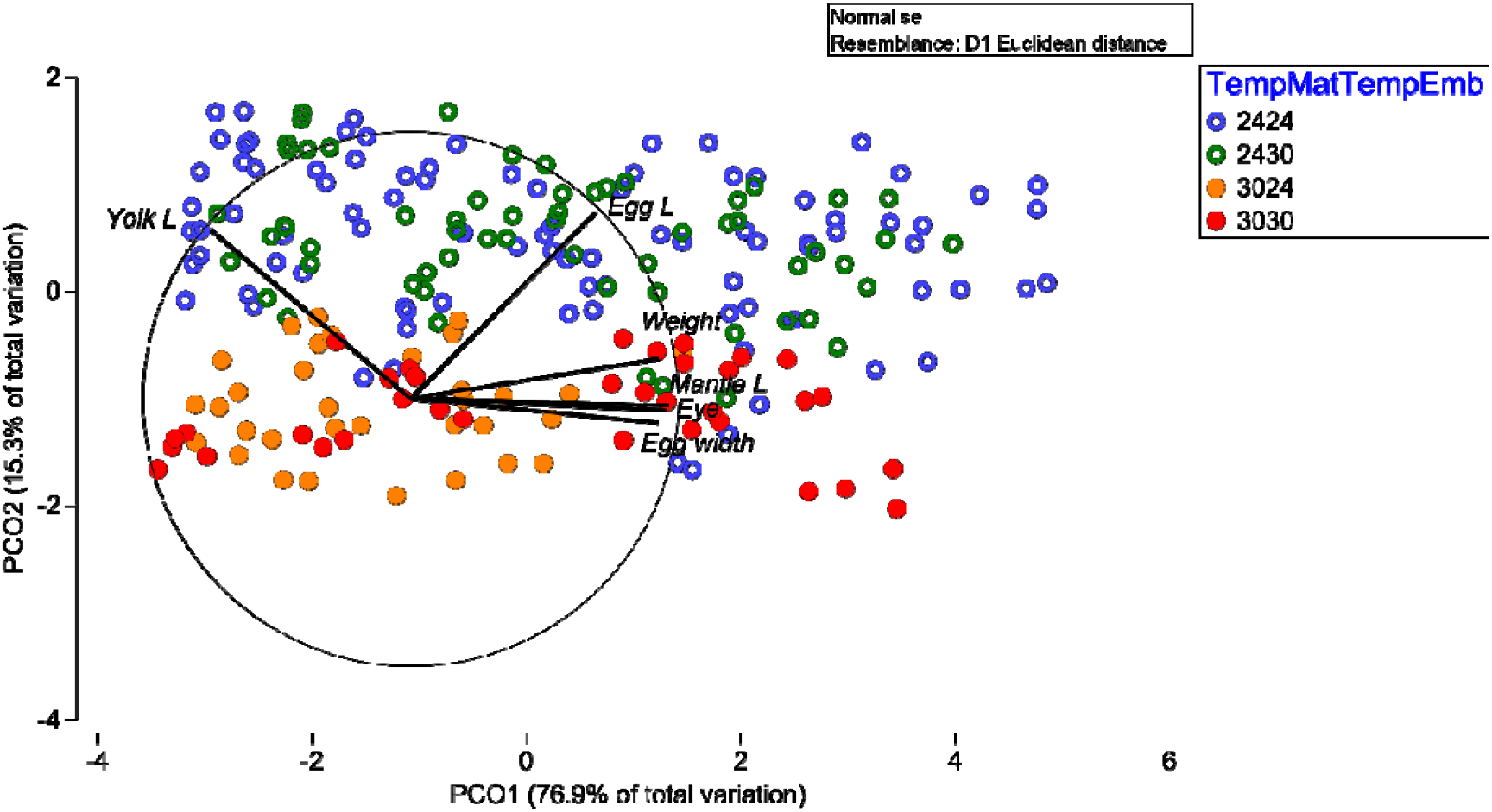
Principal coordinates (a) PCO1 vs PCO2 of the transgenerational effect of *Octopus maya* female thermal condition (24: blue and green; and 30°C: orange and red) on morphological characteristics of the embryos exposed at 24°C and 30°C. In the codes for the treatments, the first number identifies the thermal condition of females and the second one thermal condition of embryos.

Embryos from females maintained at 24°C and incubated at 24°C showed higher wet weight, mantle length, eye diameter, yolk and egg length than the embryos incubated at 30°C. In contrast, no statistical differences were observed in morphological characteristics of the embryos from females maintained at 30°C both incubated at 24 and 30°C (Fig. 3).

### 3.3 Antioxidant enzymes and oxidant damage

In the activation phase, PERMANOVA indicated that the variable maternal temperature and embryo incubation temperature have a significant effect on antioxidant defense mechanisms and oxidant damage. Pair-wise tests indicated statistical differences for pair levels of maternal temperature within levels of embryo temperature during the activation phase. The antioxidant defense mechanism of the embryos from the females maintained at 24°C and incubated at 24°C showed higher concentrations of PO and CAT activity. Organisms from females maintained at 24°C and incubated at 30°C showed high PO and CAT activities and high variability in GST and GSH values. Samples from embryos from females maintained at 24°C and incubated at 30°C showed the highest PO and GST levels. In contrast, the results of the embryos from females maintained at 30°C showed low activities or concentrations of all the tested variables, especially lower in embryos from females both maintained and incubated at 30°C.

The results of the antioxidant defense mechanisms during the growth phase indicated significant differences in the interaction of the variables maternal and incubation temperatures of embryos. The Pair-wise tests indicated statistical differences for pair levels of maternal temperature within embryo temperature levels. Samples of embryos from females kept and incubated at 30°C showed lower CAT activity, total GSH concentration and higher activities of GST. Samples from female embryos maintained at 24°C and kept at 30°C showed the highest PO and GST levels.

### 3.4. Esterase

Esterase activities were not affected by the thermal origin of the embryos nor incubation temperature (*p* □ 0.05; Fig. 5). Differences in activities of CbE and AChE were only detected according to development stage, with low values at heart activation and circulatory system activation in comparison with activities measured during the growth phase (SS 1507; DF = 1; MS 1507; F = 44.9; p □ 0.0001; Fig. 5).

**Fig. 4.**
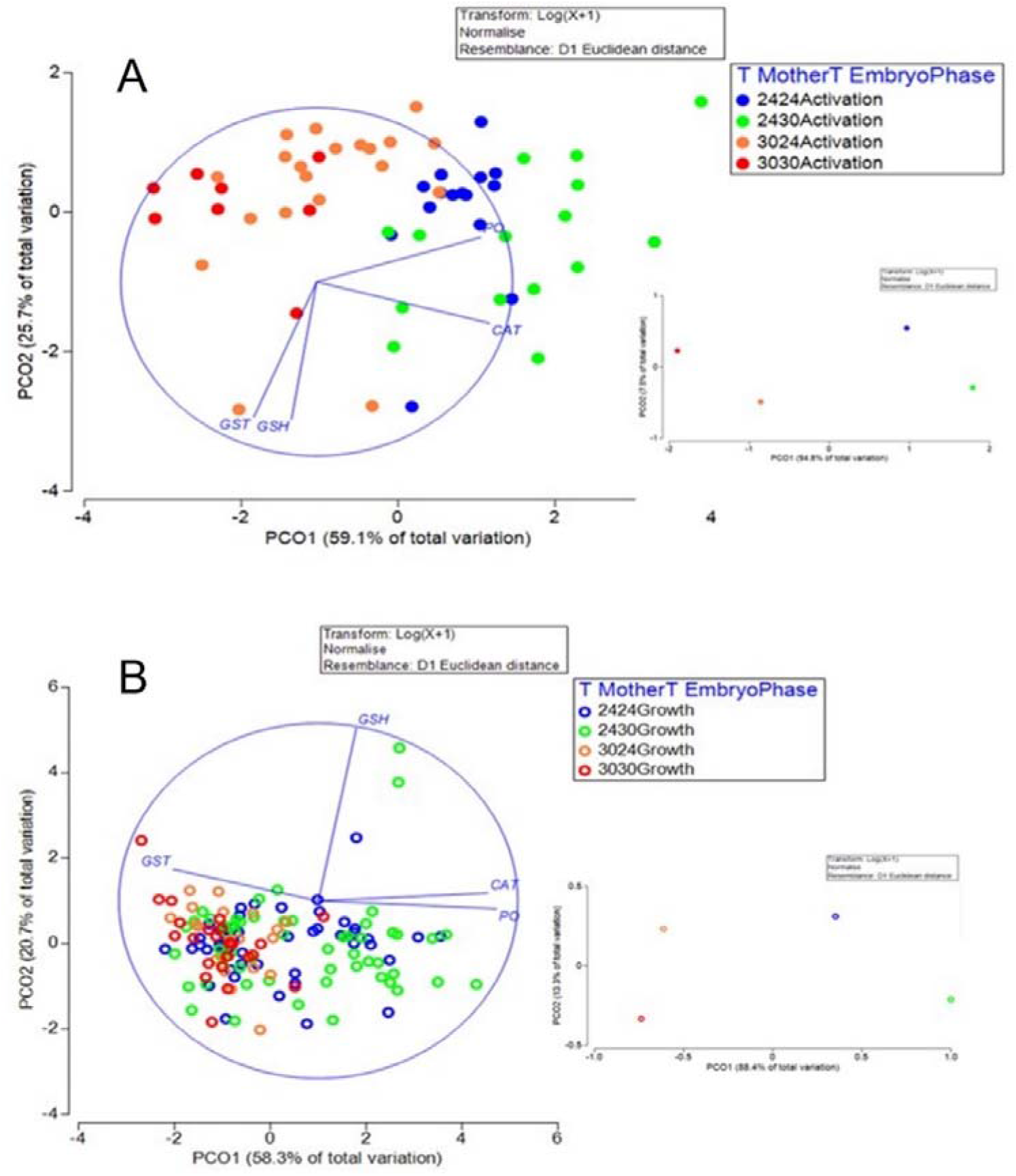
Principal coordinates PCO1 vs PCO2 of the transgenerational effect of *Octopus maya* female thermal condition (24: blue; and 30°C: red) on antioxidant defense mechanisms of the embryos at activation stage (A), and growth stage (B) and exposed at 24°C and 30°C. In the symbols, the first number identifies the thermal condition of females and the text the phase of embryo development. Centroids of each PCO at the right side of each one. CAT = Catalase, GST = Glutathione-s-Transferace, GSH = Total Glutathione, PO = Protein carbonilation.

**Fig. 5.**
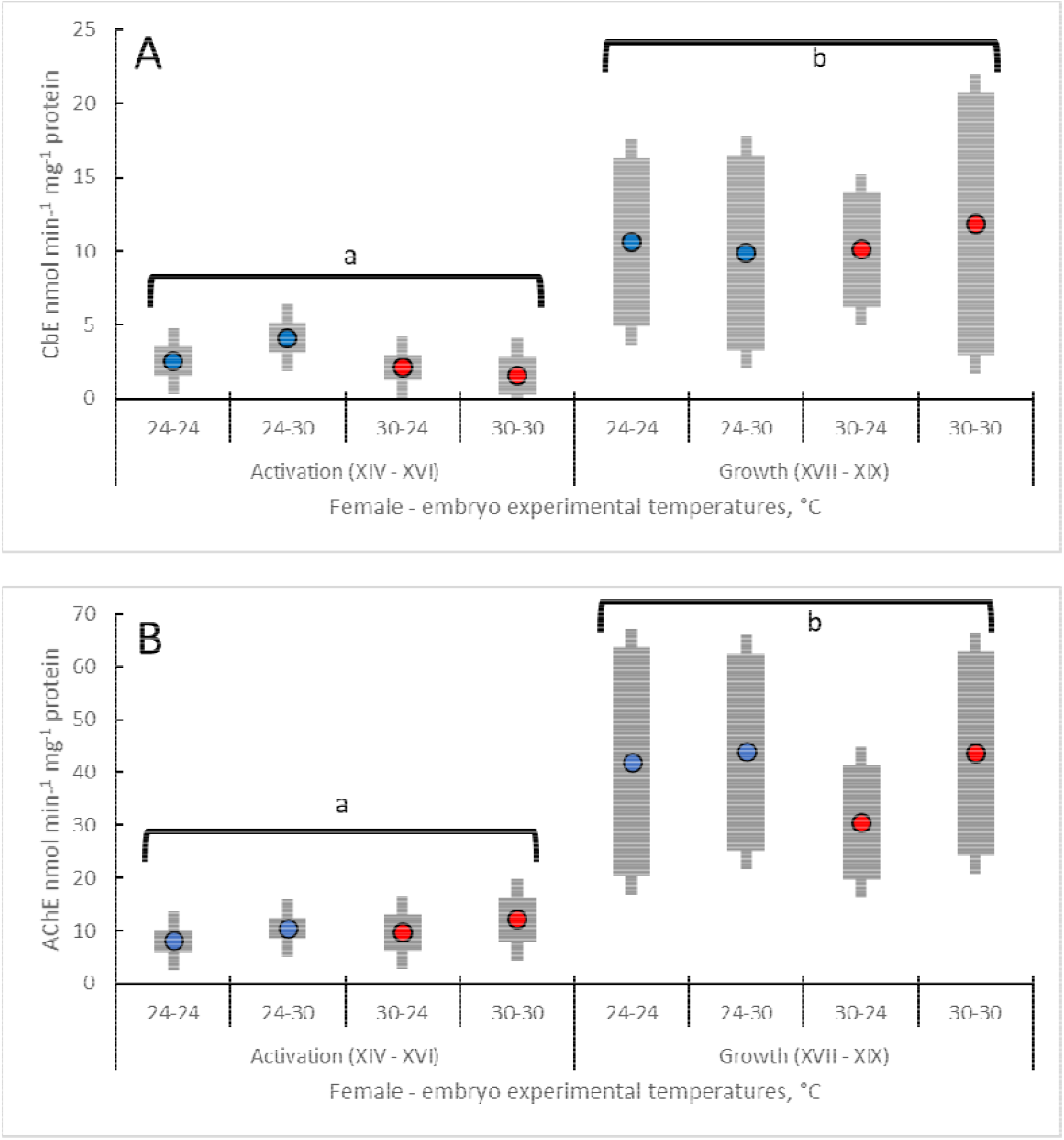
Effect of incubation temperature on Carboxylesterase (CbE) and acetylcholinesterase (AChE) activities of *Octopus maya* embryos (Stages XIV – XVI = activation; VII – XIX = growth) from females acclimated at 24°C (blue circle) and 30°C (red circle). The first number on X-axis indicates the female temperature acclimation. The second number indicates the embryo incubation temperature. Values ± SD. Different letters mean statistical differences between phases at *p* □ 0.0001.

## 4. Discussion

Several studies made in *O. maya* embryos, juveniles and adults have demonstrated that this species is relatively sensitive to high temperatures, of which the thermal limit for embryos and spawners is around 27°C (Ángeles-González et al., 2021; Caamal-Monsreal et al., 2016; Domínguez-Estrada et al., 2022; Juárez et al., 2015, 2016, 2022; López-Galindo et al., 2018; Meza-Buendia et al., 2021; Noyola et al., 2013). In a previous study, Juarez et al. observed that juveniles of thermally stressed females had a growth rate of approximately half of that recorded in animals of non-stressed females, suggesting that a transgenerational high temperature effect could be occurring in this octopus species (Juarez et al., 2016). Additionally, these authors observed that egg fecundity was affected by temperatures of 30°C (Juárez et al., 2015). The existence of endocrine control mechanisms in optic (Domínguez-Estrada et al., 2022) and oviducal glands to inhibit spawning at elevated temperatures (Juárez et al., 2022) was recently demonstrated, suggesting that when the temperature is stressing, females avoid potential stress for embryos inhibiting spawning. These results support the hypothesis assuming that in warming scenarios, adults could migrate to deeper waters searching for lower temperatures (22 to 26°C) where reproductive processes can occur successfully (Angeles-Gonzalez et al., 2021; Pimentel et al., 2012). Unfortunately, ecological adjustments as migrations require time (decades, hundreds of years, millennia?) due to deep and complex modifications in the ecosystem that is invaded and inevitably provokes overlapping niches (Ángeles-González et al., 2021; Eddy et al., 2020). Sadly, ocean warming is occurring faster than physiological adjustments in species and structure of the ecosystems, leaving adaptations linked to phenotypic plasticity of the population ability (or not) to respond to the rapid environmental changes (Tittensor et al., 2021). In this sense, it is highly probable that female octopuses that cannot migrate to cooler waters during ovarian maturation (i.e. due to niche overlap with *O. americanus* (Avendaño et al., 2020)) would experience negative transgenerational consequences of high temperatures in their progeny.

Several potential mechanisms can explain the negative transgenerational effect of thermal stress in *O. maya*. First, embryos from thermally stressed females had morphological alterations, showing a smaller amount of yolk and smaller sizes than embryos of non-stressed females. Second, embryos from thermally stressed females had higher metabolic rates than those observed in embryos of non-stressed females. Third, embryos from thermally stressed females had lower GSH levels and lower CAT activities, which indicate a higher ROS level during activation and growth developmental phases, and also that embryos from thermally stressed females could have added stress due to ROS accumulation during embryo development (Olivares et al., 2019)

The negative transgenerational effects of thermal stress in *O. maya* can be associated with the use of yolk reserves and consequences on embryo morphology and physiology, which compose an energetic mechanism. In octopus embryos, as in many other aquatic organisms, yolk quantity and quality are key aspects of this phase of their life cycle (Caamal-Monsreal et al., 2016). In fish, embryonic and early larval development and metabolism are fueled entirely by maternally-derived nutritional resources (yolk and oil) before the onset of exogenous feeding, which is a good indicator of parental physiological condition (Hou and Fuiman, 2022). Thus, the transgenerational effect of high temperature in *O. maya* could start with the inability of females to store enough yolk (or yolk with enough quality?) in the eggs. Previous experiments made at UNAM-Sisal laboratory demonstrated that adult females exposed to 30°C experienced a reduction in aerobic scope. This result suggests that in such thermal conditions, females had not enough energy to reach an adequately ovarian maturation (Meza-Buendia et al., 2021), even when in laboratory conditions, females were fed ad libitum with a paste that covered nutritional requirements for reproduction (Tercero-Iglesias et al., 2015). In this sense, hypothesizing that embryos from thermally stressed females were equipped with limited yolk quantities (quality?) by reducing the energy available for tissue and organ synthesis, the final size of the animals was affected (Fig. 6). In adults of the coral colonies of *Pocillopora acuta* exposed to thermal stress, offspring were also smaller than those obtained in adults without thermal stress, which suggests, as observed in octopus embryos, that smaller progeny can be a transgenerational effect in other ectotherm animals (McRae et al., 2021).

**Fig. 6.**
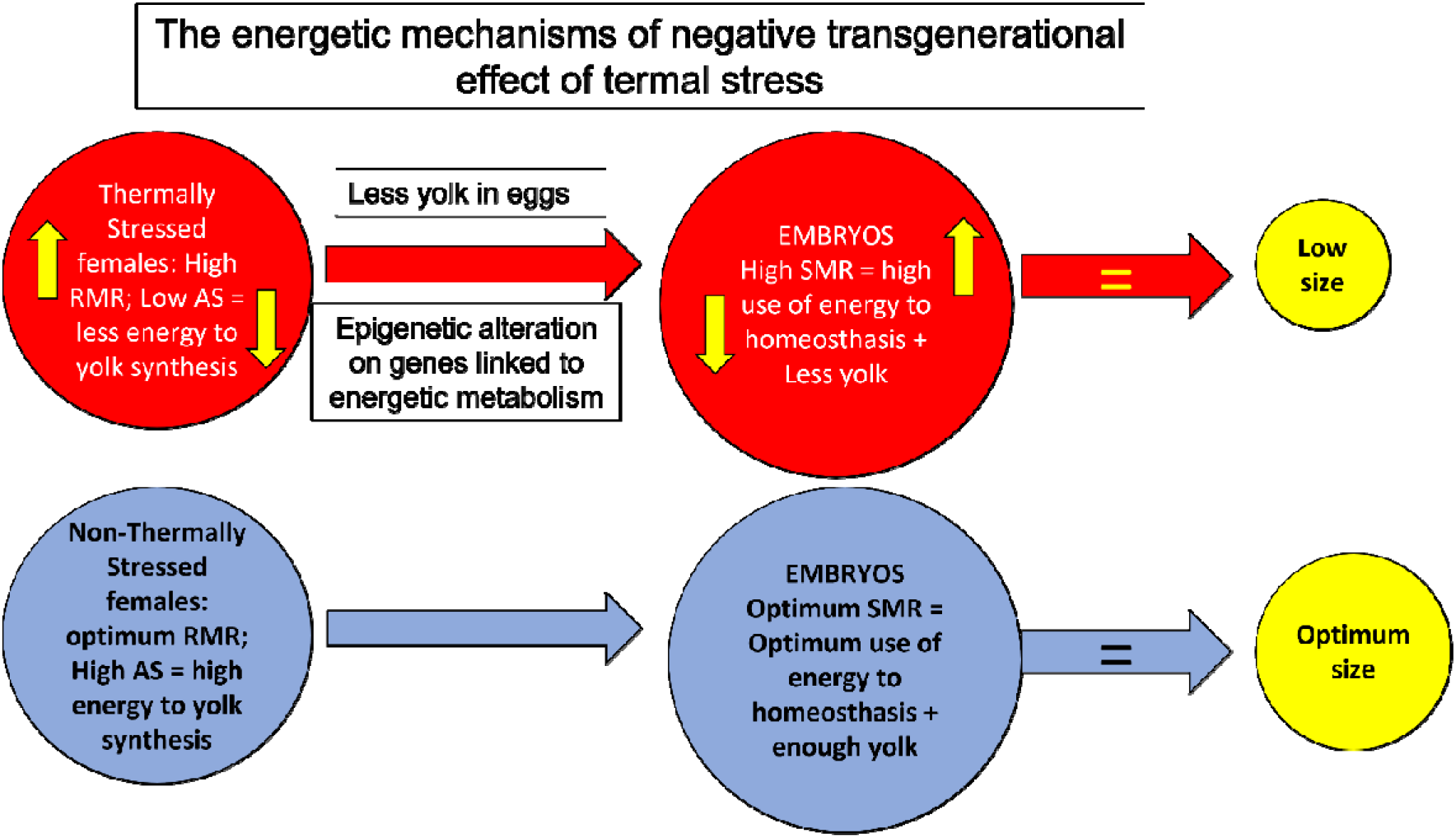
The flow diagram shows the hypothesis to explain the negative transgenerational effect of maternal thermal stress in *O. maya* embryos (red circle) and embryos from non-stressed females (blue circle). Yellow arrows indicate that although thermally stressed females had a high routine metabolic rate, this stress prevents the use of enough energy for yolk synthesis. Also, epigenetic alterations could modify the energetic metabolism, provoking high metabolic rates in embryos. As a result, embryos from thermally stressed females with high metabolic rates have less yolk and, consequently, a smaller size than embryos laid by non-stressed females.

Although the reason why embryos of thermally stressed females have high metabolic rates is still not known, we may hypothesize that some epigenetic alterations could be involved (Eirin-Lopez and Putnam, 2019; Zhao et al., 2018) (Fig. 6). Invertebrates have wide breathing strategies to facilitate oxygen extraction of the surrounding water (Bell and Syed, 2012). In cephalopod embryos, in the absence of any circulatory mechanism to aid oxygen transport to the tissue, oxygen must pass by diffusion from the external environment through the egg capsule to the embryo (Pimentel et al., 2012). During embryo development, changes occur in surface area and reduction of the egg wall thickness when, at the same time, the metabolic demand rises due to cellular growth, organogenesis, and muscular activity (Caamal-Monsreal et al., 2016; Cronin and Seymour, 2000; Sánchez-García et al., 2017; Uriarte et al., 2012). What kind of mechanisms could then be epigenetically altered because of parental thermal stress?

According to Eirin-López and Putnam, (2019), a wide range of epigenetic mechanisms might be modulated because of environmental stress, ending up with changes in the phenotypes of organisms. One of these stressors is the temperature, which is known to modify the use of nutrients that are channeled to yolk production (Caamal-Monsreal et al., 2016). It has been reported that levels of acetyl-CoA, a donor to acetylation reactions, are correlated with the extent of histone acetylation and, that this epigenetic mark favors gene expression (Gibney and Nolan, 2010; Lu and Thompson, 2012; Sidoli et al., 2019; Tsuchiya et al., 2014). If embryos from stressed females have less yolk content, it can be proposed that suboptimal available levels of acetyl-CoA might thus trigger a histone hypoacetylation event which in turn might favor the silencing of a set of genes (probably, metabolism-related genes) that under normal embryo development conditions are necessary for growth. In fact, a direct link between the acetyl-CoA levels and the subsequent acetylation of histones at growth-related genes has been observed (Cai et al., 2011). Another mechanism by which hypoacetylation might be occurring is because elevated temperatures could alter the degradation of amino acids (AA) in the female during yolk synthesis processes in the embryo. If temperature modifies how AA are used as a cellular energy source in the ovary, then NAD+ can be synthesized *de novo* from AA as tryptophan. NAD+ is an obligatory co-factor of sirtuin (SIRT1 and SIRT 6) activities, which deacetylate histones H3K9/14 and H3K9/56, respectively. The deacetylation of histone H3 by these SIRT could down-regulate the expression of metabolic genes in the embryo, thereby altering various metabolic pathways. The latter would cause uncoupling blocks 1 and 2, (the latter comprising mostly the tri-carboxylic acid cycle) with block 3 (electron transport system ETS); the consequent escape of protons and increase in ROS production (Etchegaray and Mostoslavsky, 2016) provokes at the same time an increment in mitochondrial oxygen consumption as a compensatory reaction (Wallace and Fan, 2010), which is consistent with what was observed in embryos of *O. maya* (Fig. 2). Concurrently, several questions can be addressed. Is it possible that as a consequence of epigenetic alteration, octopus embryos from thermally stressed females have more mitochondria than those coming from non-stressed females?. Are more enzymes participating in energetic pathways of the embryos even if they have the same number of mitochondria? Do embryos have a lower chorion thickness that facilitates oxygen flow from water to the embryo? Moreover, other possibilities could be included in the epigenetic alterations due to parental thermal stress. Imperadore et al. (2019) showed that two pallial nerves innervate two stellate ganglia in *O. vulgaris* mantle, controlling mantle breathing movements. Are there any chances that epigenetic alterations affect the embryo nervous central system changing the control ability of the pallial nerves and provoking an uncontrolled muscle activity in embryos during activation and growth phases, enhancing energy requirements and causing higher oxygen consumption? AChE are β-esterases that degrade choline-based esters; their role in cephalopods, as in many other organisms, is to act as a neurotransmitter in the nervous system, controlling excitatory stimulus (Omedes et al., 2022). This study did not observe that female thermal stress provoked alterations in AChE activity levels, suggesting that if thermal stress provoked alterations in the nervous system, it was not through AChE inhibition. However, AChE and other esterases have other physiological roles, such as cell differentiation, apoptosis and arm regeneration, suggesting that other aspects of the embryo metabolism should be studied to find how the epigenome could be altering *O. maya* embryo metabolism.

Among the mechanisms involved in maintaining homeostasis in aquatic invertebrates, antioxidant defense mechanisms (ANTIOX) have a key role in neutralizing ROS and RNS overproduction (Abele et al., 2007; Rodríguez-Fuentes et al., 2017). Although ANTIOX mechanisms work all the time to neutralize ROS, in extreme temperatures, the rate of ROS production overwhelms ANTIOX capability and repair mechanisms, leading to cellular disparities (Rahaman and Rahaman, 2021). These systems might sometimes fail to counteract the toxic effects of an amplified oxygen and nitrogen radical formation, elevating lipid peroxidation levels and producing cellular oxidative stress (Regoli and Giuliani, 2014). The magnitude of oxidative stress depends on temperature and exposure time besides the ability of organisms to upregulate the enzymes involved in antioxidative defenses (Matozzo et al., 2013; Parisi et al., 2021). In this study, changes in the activities of antioxidant enzymes and GSH levels indicate the ability of embryos to deal with ROS and oxidative damage (measured as PO). In many animals, the most abundant cytosolic scavenger is the reduced glutathione (GSH), a tri-peptide (g-glutamyl-cysteinyl glycine), which directly neutralizes several reactive species through its oxidation to glutathione disulfide (GSSG). In addition, GSH acts as a cofactor of several antioxidant glutathione-dependent enzymes reducing many ROS (H_2_O_2_, O_2_^-^, HO^+^ and lipid hydroperoxides) with high efficiency (Rahaman and Rahaman, 2021) (Regoli and Giuliani, 2014); its depletion during oxidative stress is related to an intensive use as either first-line antioxidant or as an enzyme cofactor. It is interesting to note that carbonyl groups in oxidized proteins (PO) and CAT activity were higher in embryos from non-thermally stressed females than those from stressed females suggesting differences in the form in which ROS and ANTIOX acted in the two groups of embryos. In *O. mimus* embryos, a part of ROS and oxidized biomolecules are transferred by the female through the yolk even in optimal thermal conditions (Olivares et al., 2019). To neutralize the ROS inherited and the ones produced at respiration, embryos turn on the antioxidant mechanisms once the circulatory system is activated, allowing the enzymes to neutralize ROS after organogenesis when the heart and circulatory system start working (Sánchez-García et al., 2017). A similar sequence of events was observed in *O. maya* embryos of non-stressed females. High levels of PO at the time of activation may be closely related to baseline levels plus PO produced during respiration, which is more evident in thermally stressed embryos where CAT activity also increases. In the case of embryos of thermally stressed females, other mechanisms could be operating. In those embryos, it is evident that the antioxidant system is collapsing, CAT activities and GSH concentrations are lower, and GST activity increases, especially at the growth phase. GST is an enzyme that has two important roles; both use GSH. First as a detoxification enzyme, conjugating by-products with GSH and taking them out of the cell, and second, stopping the chain reaction of lipid peroxidation (Regoli, 2014). PO concentrations may be relatively lower in the non-thermally stressed females, but it may only indicate that PO was aggregated, no longer soluble (as in healthy cells) and unmeasurable with the method used, as it has been previously described (Maisonneuve et al., 2008).

## 5. Conclusion

The results in this research are consistent with the adverse parental carry-over of thermal effects, which may be associated with trade-offs between the female condition (less aerobic scope and energy for yolk synthesis at high temperatures) and the investment required (yolk quality/quantity?) to have a successful offspring. Embryos from thermally stressed females had smaller sizes, less yolk, and higher metabolic rates. Additionally, a collapse in the antioxidant defense system was observed in embryos from thermally stressed females, indicating they were unable to control the high load of ROS and oxidative damage, which was partially acquired by maternal inheritance.

To conclude, further research should (i) assess if, as a consequence of epigenetic alteration, octopus embryos -from thermally stressed females-have more mitochondria than those coming from non-stressed females; (ii) confirm, if more enzymes are participating in the embryo energetic pathways even if they have the same number of mitochondria; (iii) observe if the embryo chorion wall thickness depends on the female thermal condition. Confirm if global acetylations levels are minors in embryos coming from stressed females. Other possibilities could be included in epigenetic alterations due to parental thermal stress. Thus, future studies need be performed with the purpose of knowing if epigenetic alterations affect the organism nervous central system, changing the control ability of pallial nerves and provoking an uncontrolled muscle activity during activation and growth phases, enhancing energy requirements, and causing higher oxygen consumption

## 6. Acknowledgments and funding

CR thanks Consejo Nacional de Ciencia y Tecnología for funding the project CONACYT 61503 and PAPIIT IN203022; D. Fischer provided English Edition.

## Literature Cited

Abele, E., Philip, E., Gonzalez, P.M., Puntarulo, S., 2007. Marine invertebrate mitochondria and oxidative stress. Front. Biosci. 12, 933–946.

Angeles-Gonzalez, L.E., Calva, R., Santos-Valencia, J., Avila-Poveda, O.H., Olivares, A., Díaz, F., Rosas, C., 2017. Temperature modulates spatio-temporal variability of the functional reproductive maturation of Octopus maya (Cephalopoda) on the shelf of the Yucatan Peninsula, Mexico. Journal of Molluscan Studies in press: doi:10.1093/mollus/eyx013, 1–9.

Ángeles-González, L.E., Lima, F.D., Caamal-Monsreal, C., Díaz, F., Rosas, C., 2020. Exploring the effects of warming seas by using the optimal and pejus temperatures of the embryo of three Octopoda species in the Gulf of Mexico. J Therm Biol 94, 102753. Doi: 102710.101016/j.jtherbio.102020.102753.

Ángeles-González, L.E., Martínez-Meyer, E., Rosas, C., Guarneros-Narváez, V., López-Rocha, J., Escamilla-Ake, A., Osorio-Olvera, L., Yáñez-Arenas, C., 2021. Long term environmental data explain better 1 the abundance of the red octopus (Octopus maya) when testing the niche centroid hypothesis. J Exp Mar Biol Ecol in press.

Avendaño, O., Roura, A., Cedillo-Robles, C.E., Gonzalez, A., Rodrıguez-Canul, R., Velazquez-Abunader, I., Guerra, A., 2020. Octopus americanus: a cryptic species of the O. vulgaris species complex redescribed from the Caribbean. Aquat Ecol 54, 909 –925, Doi: 910.1007/s10452-10020-09778-10456.

Bell, H.J., Syed, N.I., 2012. Control of breathing in invertebrate model system. Comprehensive Physiology 2, 1745–1766.

Byrne, M., Foo, S.A., Ross, P.M., Putnam, H.M., 2019. Limitations of cross-and multigenerational plasticity for marine invertebrates faced with global climate change. Glob. Change Biol 26, 80 -102. Doi: 110.1111/gcb.14882.

Caamal-Monsreal, C., Uriarte, I., Farias, A., Díaz, F., Sánchez, A., Re, A.D., Rosas, C., 2016. Effects of temperature on embryo development and metabolism of O. maya. Aquaculture 451, 156–162.

Cai, L., Sutter, B.M., Li, B., Tu, B.P. 2011. Acetyl-CoA induces cell growth and proliferation by promoting the acetylation of histones at growth genes. Mol Cell 42(4), 426–437.

Cronin, E.R., Seymour, R.S., 2000. Respiration of the eggs of the giant cuttlefish Sepia apama. Mar.Biol. 136, 863–870.

Dominguez-Castanedo, O; Palomino-Cruz, D; Mascaró, M.; Rodríguez-Fuentes, G; Juárez, O.E.; Galindo-Sánchez, C.E.; Caamal-Monsreal, C.; Galindo-Torres, P.; Díaz, F.; Rosas, C. 2022. Data of transgenerational effects of thermal stress in embryos of O. maya. Zenodo Doi: 10.5281/zenodo.6533870

Domínguez-Estrada, A., Galindo-Sánchez, C., Ventura-López, C., Rosas, C., Juárez, O., 2022. Optic gland pathways in response to thermal stress through the reproductive phase of Octopus maya females. Journal of Molluscan Studies In press.

Eddy, T.D., Bernhardt, J.R., Blanchard, J.L., Cheung, W.L., Colléter, M., du Pontavice, H., Fulton, E.A., Gascuel, D., Kearney, K.A., Petrik, C. M.,, Roy, T., Rykaczewski, R.R., Selden, R., Stock, C.A., Wabnitz, C.C., Watson, R.A., 2020. Energy Flow Through Marine Ecosystems: Confronting Transfer Efficiency. Trends in Ecology & Evolution 36, 76–86. Doi: 10.1016/j.tree.2020.1009.1006.

Eirin-Lopez, J.M., Putnam, H.M., 2019. Marine Environmental Epigenetics. Annu. Rev. Mar. Sci. 11, 335–368, Doi: 310.1146/annurev-marine-010318-095114.

Etchegaray, J.P., Mostoslavsky, R., 2016. Interplay between Metabolism and Epigenetics: A Nuclear Adaptation to Environmental Changes. Molecular Cell 62, 695–711.

Feidantsis, K., Georgoulis, J., Giantsis, I., Michaelidis, 2021. Treatment with ascorbic acid normalizes the aerobic capacity, antioxidant defence, and cell death pathways in thermally stressed Mytilus galloprovincialis. Comparative Biochemistry and Physiology, Part B 255

Fellous, A., Favrel, P., Riviere, G., 2015. Temperature influences histone methylation and mRNA expression of the Jmj-C histone-demethylase orthologues during the early development of the oyster Crassostrea gigas. Marine Genomics 19, 23–30. Doi: 10.1016/j.margen.2014.1009.1002.

Fellous, A., Wegner, K.M., John, U., Mark, F.C., 2021. Windows of opportunity: Ocean warming shapes temperature-sensitive epigenetic reprogramming and gene expression across gametogenesis and embryogenesis in marine stickleback. Glob. Change Biol 28, 54–71. Doi: 10.1111/gcb.15942

Fridovich, I., 1986. Biological effects of the superoxide radical. Arch Biochem Biophys 247, 1–11, Doi: 10.1016/0003-9861(1086)90526-90526.

Gibney, E.R., Nolan, C.M. 2010. Epigenetics and gene expression. Heredity 105, 4– 13.

Hayward, L.S., Wingfield, J.C., 2004. Maternal corticosterone is transferred to avian yolk and may alter offspring growth and adult phenotype.. Gen.Comp.Endocrinol. 135: 365–371.

Hou, Z., Fuiman, L.A., 2022. Incorporation of dietary lipids and fatty acids into red drum Sciaenops ocellatus eggs. Com.Biochem.Physiol. B 258, 110694. Doi: 110610.111016/j.cbpb.112021.110694

Juárez, O., Arreola-Meráz, L., Sánchez-Castrejón, E., Avila-Poveda, O.H., López-Galindo, L., Rosas, C., Galindo-Sánchez, C., 2022. Oviducal gland transcriptomics of Octopus maya through physiological stages and the negative effects of temperature on fertilization. PeerJ 10, e12895. Doi: 12810.17717/peerj.12895.

Juárez, O., Galindo, C.E., Díaz, F., Re, A.D., Sanchez-García, A.M., Caamal-Monsreal, C., Rosas, C., 2015. Is temperature conditioning Octopus maya fitness? J. Exp. Mar. Biol. Ecol. 467, 71–76.

Juarez, O., Hau, V., Caamal-Monsreal, C., Galindo, C.E., Díaz, F., Re, A.D., Rosas, C., 2016. Effect of maternal temperature stress before before spawning over the energetic balance of Octopus maya juveniles exposed to a gradual temperature changes. J. Exp. Mar. Biol. Ecol. 474, 39–45.

Juárez, O., López-Galindo, L., Pérez-Carrasco, L., Lago-Lestón, A., Rosas, C., Di Cosmo, A., Galindo-Sánchez, C., 2019. Octopus maya white body show sexspecific transcriptomic profiles during the reproductive phase, with high differentiation in signaling pathways. PloS ONE 14, e0216982. https://doi.org/0216910.0211371/journal.pone.0216982.

Klosing, A., Casas, E., Hidalgo-Carcedo, C., Vavouri, T., Lehner, B., 2019. Transgenerational transmission of environmental information in C. elegans. Science 356, 320-323. Doi: 310.1126/science.aah6412.

Lin, D., Han, F., Xuan, S., Chen, X., 2019. Fatty acid composition and the evidence for mixed income–capital breeding in female Argentinean short<sub>lZI</sub> fin squid Illex argentinus. Mar. Biol. 166, 90. DOI: 10.1007/s00227-00019-03534-00220.

López-Galindo, L., Galindo-Sánchez, C., Olivares, A., Avila-Poveda, O.H., Díaz, F., Juárez, O.E., Lafarga, F., Pantoja-Péreza, J., Caamal-Monsreal, C., Rosas, C., 2018. Reproductive performance of Octopus maya males conditioned by thermal stress. Ecological Indicators 96 437–447 doi: 410.1016/j.ecolind.2018.1009.1036.

Lu, C., Thompson, C.B. 2012. Metabolic regulation nof epigenetics. Cell Metab 16(1), 9–17.

Maisonneuve, E., Fraysse, L., Lignon, S., Capron, L., Dukan, S. 2008 Carbonylated proteins are detectable only in a degradation resistant aggregate state in Escherichia coli. J Bacteriol 190,20 6609–6614 doi:10.1128/JB.00588-08

Matozzo, V., Chinellato, A., Munari, M., Bressan, M., Marin, M.G., 2013. Can the combination of decreased pH and increased temperature values induce oxidative stress in the clam Chamelea gallina and the mussel Mytilus galloprovincialis? Mar Pollut Bull 72, 34–40, Doi: 10.1016/j.marpolbul.2013.1005.1004.

McRae, C., Huang, W.-B., Fan, T.-Y., Cóté, I.M., 2021. Effects of thermal conditioning on the performance of Pocillopora acuta adult coral and their offspring. Coral reefs 40, 1491–1503. doi: 1410.1007/s00338-00021-02123-00339.

Meza-Buendia, A.K., Trejo-Escamilla, I., Piu, M., Caamal-Monsreal, C., Rodríguez-Fuentes, G., Díaz, F., Re, A.D., Galindo-Sánchez, C.E., Rosas, C., 2021. Why high temperatures limit reproduction in cephalopods? The case of Octopus maya. Aquacult. Res. DOI: 10.1111/are.15387.

Naef, A., 1928. Die Cephalopoden (Embryologie) Fauna e flora del Golfo di Napoli 35, 1–375.

Noyola, J., Caamal-Monsreal, C., Díaz, F., Re, A.D., Sánchez, A., Rosas, C., 2013. Thermal preference, tolerance and metabolic rate of early juveniles of Octopus maya exposed to different acclimation temperatures. J. Therm. Biol. 38 14–19.

Olivares, A., Rodríguez-Fuentes, G., Mascaró, M., Sánchez, A., Ortega, K., Caamal-Monsreal, C., Tremblay, N., Rosas, C., 2019. Maturation trade-offs in octopus females and their progeny: energy, digestion and defence indicators. PeerJ 7, e6618. Doi: 6610.7717/peerj.6618.

Omedes, S., Andrade, M., Escolar, O., Villanueva, R., Freitas, R., Solé, M., 2022. B-esterases characterisation in the digestive tract of the common octopus and the European cuttlefish and their in vitro responses to contaminants of environmental concern. Environmental Research 210, 112961 Doi: 112910.111016/j.envres.112022.112961.

Parisi, M.G., Giacoletti, C., Mandaglio, M., Sará, G., 2021. The entangled multi-level responses of Mytilus galloprovincialis (Lamarck, 1819) to environmental stressors as detected by an integrated approach. Marine Environmental Research 168.

Pimentel, M.S., Trübenbach, K., Faleiro, F., Boavida-Portugal, J., Repolho, T., Rosa, R., 2012. Impact of ocean warming on the early ontogeny of cephalopods: a metabolic approach. Mar Biol 159, 2051–2059, doi: 2010.1007/s00227-00012-01991-00229.

Pörtner, H.O., 2010. Oxygen-and capacity-limitation of thermal tolerance: a matrix for integrating climate-related stressor effects in marine ecosystems. J. Exp. Biol. 213, 881–893.

Pörtner, H.O., Bock, C., Mark, F.C., 2017. Oxygen-and capacity-limited thermal tolerance: bridging ecology and physiology. Journal of Experimental Biology 220, 2685 –2696, doi:2610.1242/jeb.134585.

Pörtner, H.O., Roberts, D.C., Poloczanska, E.S., Mintenbeck, K., Tignor, M., Alegría, A., Craig, M., Langsdorf, S., Löschke, S., Möller, V., 2022. Climate Change 2022: Impacts, Adaptation, and Vulnerability. Contribution of Working Group II to the Sixth Assessment Report of the Intergovernmental Panel on Climate Change in: Pörtner, H.O., Roberts, D.C., Tignor, M., Poloczanska, E.S., Mintenbeck, K., Alegría, A., Craig, M., Langsdorf, S., Löschke, S., Möller, V., A., O., Rama, B. (Eds.), IPCC, 2022: Summary for Policymakers Cambridge University Press., Cambridge.

Rahaman, M.S., Rahaman, M.S., 2021. Effects of elevated temperature on prooxidant-antioxidant homeostasis and redox status in the American oyster: Signaling pathways of cellular apoptosis during heat stress. Environmental Research 196 Doi: 10.1016/j.envres.2020.110428.

Regoli, F., Giuliani, M.E., 2014. Oxidative pathways of chemical toxicity and oxidative stress biomarkers in marine organisms. Marine Environmental Research 93, 106–117.

Repolho, T., Baptista, M., Pimentel, M.S., Dionısio, G., Trübenbach, K., Lopes, V.M., Lopes, A.R., Calado, R., Diniz, M., Rosa, R., 2014. Developmental and physiological challenges of octopus (Octopus vulgaris) early life stages under ocean warming. J. Comp. Physiol. B 184, 55–64.

Rodríguez-Fuentes, G., Murúa-Castillo, M., Díaz, F., Rosas, C., Caamal-Monsreal, C., Sánchez, A., Paschke, K., Pascual, C., 2017. Ecophysiological biomarkers defining the thermal biology of the Caribbean lobster Panulirus argus. Ecological Indicators in press, doi: 10.1016/j.ecolind.2017.03.011.

Roumbedakis, K., Mascaró, M., Martins, M.L., Gallardo, P., Rosas, C., Pascual, C., 2017. Health status of post-spawning Octopus maya (Cephalopoda: Octopodidae) females from Yucatan Peninsula, Mexico. Hydrobiologia doi: 10.1007/s10750-017-3340-y.

Sánchez-García, A., Rodríguez-Fuentes, G., Díaz, F., Galindo-Sánchez, C., Ortega, K., Mascaró, M., López, E., Caamal-Monsreal, C., Juárez, O., Noreña-Barroso, E., Re, D., Rosas, C., 2017. Thermal sensitivity of O. maya embryos as a tool for monitoring the effects of environmental warming in the Southern of Gulf of Mexico. Ecological Indicators 72, 574–585.

Secretaría de Agricultura, Ganadería, Desarrollo Rural, Pesca y Alimentación, 2014. Fishery Management Plan for octopus (O. Maya and O. Vulgaris) in the Gulf of Mexico and the Caribbean Sea. Plan-de-Manejo-Pesquero-de-Pulpo.pdf (inapesca.gob.mx).

Sidoli, S., Trefely, S., Garcia, B.A., Carrer, A. 2019. Integrated analysis of Acetyl-CoA and histone modification via mass spectrometry to investigate metabolically driven acetylation. Methods Mol Biol 1928, 125–147.

Tercero-Iglesias, J.F., Rosas, C., Mascaró, M., Poot-López, G.R., Domingues, P., Noreña, E., Caamal-Monsreal, C., Pascual, C., Estefanell, J., Gallardo, P., 2015. Effects of parental diets supplemented with different lipid sources on Octopus maya embryo and hatching quality. Aquaculture 448, 234–242.

Tsuchiya, Y., Pham, U., Hu, W., Ohnuma, S.I., Gout, I. 2014. Changes in acetyl levels during the early embryonic development of Xenopus laevis. PloS ONE 9(5): e97693. doi:10.1371/journal.pone.0097693

Uriarte, I., Espinoza, V., Herrera, M., Zuñiga, O., Olivares, A., Carbonell, P., Pino, S., Farias, A., Rosas, C., 2012. Effect of temperature on embryonic development of Octopus mimus under controlled conditions. J. Exp. Mar. Biol. Ecol. 416-417, 168–175.

Vagnerová, K., Vacková, Z., Klusonová, P., Staud, F., Kopecky, M., Ergang, P., Milkcik, I., Pácha, J., 2008. Reciprocal changes in maternal and fetal metabolism of corticosterone in rat during gestation. Reproductive Sciences 15 921–931.

Ventura-López, C., López-Galindo, L., Rosas, C., Sánchez-Castrejón, E., Galido-Torres, P., Pascual, C., Rodríguez-Fuentes, G., Juárez, O., Galindo-Sánchez, C., 2022. Sex-specific role of the optical gland in Octopus maya: a transcriptomic analysis. Gen Comp Endocrinol 320, 114000. Doi: 114010.111016/jygcen.112022.114000.

Waite, H.R., Sorte, C.J.B., 2022. Negative carry-over effects on larval thermal tolerances across a natural thermal gradient. Ecology 103, e03565. Doi:03510.01002/ecy.03565.

Wallace, D.C., Fan, W., 2010. Energetics, epigenetics, mitochondrial genetics. Mitocondrion 10, 12–31. Doi: 10.1016/j.mito.2009.1009.1006.

Zhao, L., Yang, F., Milano, S., Han, T., Walliser, E.O., Schöne, B.R., 2018. Transgenerational acclimation to seawater acidification in the Manila clam Ruditapes philippinarum: Preferential uptake of metabolic carbon. Science of the Total Environment 627, 95–10. Doi: 10.1016/j.scitotenv.2018.1001.1225.

